# Highly-parallel production of designer organoids by mosaic patterning of progenitors

**DOI:** 10.1101/2023.10.25.564017

**Authors:** Catherine M. Porter, Grace C. Qian, Samuel H. Grindel, Alex J. Hughes

## Abstract

Human organoids are a promising approach for disease modeling and regenerative medicine. However, organoid variability and limited control over morphological outcomes remain significant challenges. Here we extend a DNA ‘velcro’ cell patterning approach, precisely controlling the number and ratio of human stem cell-derived progenitors contributing to nephron and mosaic nephron/ureteric bud organoids within arrays of microwells. We demonstrate long-term control over organoid size and morphology, decoupled from geometric constraints.

## Main

Kidney organoids derived from human induced pluripotent stem cells (hiPSCs) offer a means of modeling kidney development and disease, conducting drug and genetic screens, and producing renal replacement tissue^1–5^. However, there are several roadblocks to their application, including limited organoid production scale, reproducibility, and physiologic structure, connectivity, and functionality^6^.

Previous mitigation approaches have focused on controlling biochemical signaling^2,7–9^. However, their modest success motivates new modes of engineering control over other factors, such as initial cell population size, multi-progenitor composition, and boundary conditions^9–11^, with advantages recently emerging^1,4,5,11,12^. However, scalable suspension culture methods that miniaturize organoids lack control over initial cell quantities, imparting variability in micro-organoids^4,12^, whereas microwell systems rely on geometric confinement to produce single organoids per well^13,14^. Thus, no engineering approach has combined precise control of initial size and cell composition (independent of physical boundary conditions), extended culture/imaging, and high throughput.

To decouple organoid size from boundary constraints, we adapted a high-precision, rapid cell patterning technology–photolithographic DNA-Programmed Assembly of Cells (pDPAC)^15,16^–to a microwell format suited to long-term organoid culture (**Fig. 1A**). We targeted three design opportunities: 1) precise hiPSC-derived progenitor cell number and compositional control, 2) transition of 2D patterns to self-organized 3D spheroids open to continued differentiation, and 3) sequestering individual organoids in optically-accessible microwells through 15+ day culture periods, enabling imaging and preventing aggregation (**Fig. S1**). A key advance was to follow transient 2D cell patterning on the culture substrate with a transition to 3D culture/differentiation. 2D patterning was achieved via single-stranded DNA (ssDNA) photolithographically bound with high spatial precision to a photoactive polyacrylamide (PPA) substrate on a glass slide (**Fig. S2**). Complementary lipid-conjugated ssDNA was passively incorporated into and displayed on cell membranes^15,16^. Base-pairing between cell- and substrate-bound ssDNAs thereby created temporary adhesions for cell patterning. Multiple orthogonal ssDNA sequences could be serially patterned for multiplexing cell populations. DNA micropatterns were registered and adhered to passivated, conical PDMS microwell arrays within 15.7 µm + 10.3 µm (± S.D., *n* = 16) precision (**SI Note 1, Fig. S3**). During preliminary validation, we found that microwell arrays did not interfere with Madin-Darby canine kidney cell (MDCK) patterning, retained acceptable non-specific cell adhesion properties, and minimized subsequent spheroid spreading/migration (**Fig. S4, SI Note 2, Movie S1**). pDPAC increased precision in patterned cell number for a 200-µm spot size from 12% to 4.5% coefficient of variation relative to that predicted by Poisson loading (**Fig. S4B**). After patterning, MDCKs formed 3D spheroids spontaneously within ∼6 hours, with 2% Matrigel increasing aggregation efficiency. Growth curves showed that spheroid size was predicted by ssDNA pattern diameter 1-3 days after the transition to 3D culture (**Fig. S4D**). When orthogonal ssDNA strands were used to pattern independent MDCK populations, cell number ratio accurately reflected pattern area ratio, which was retained in spheroid composition 72 hrs after the 3D transition (**Fig. S5, Movie S2**).

**Fig. 1:**
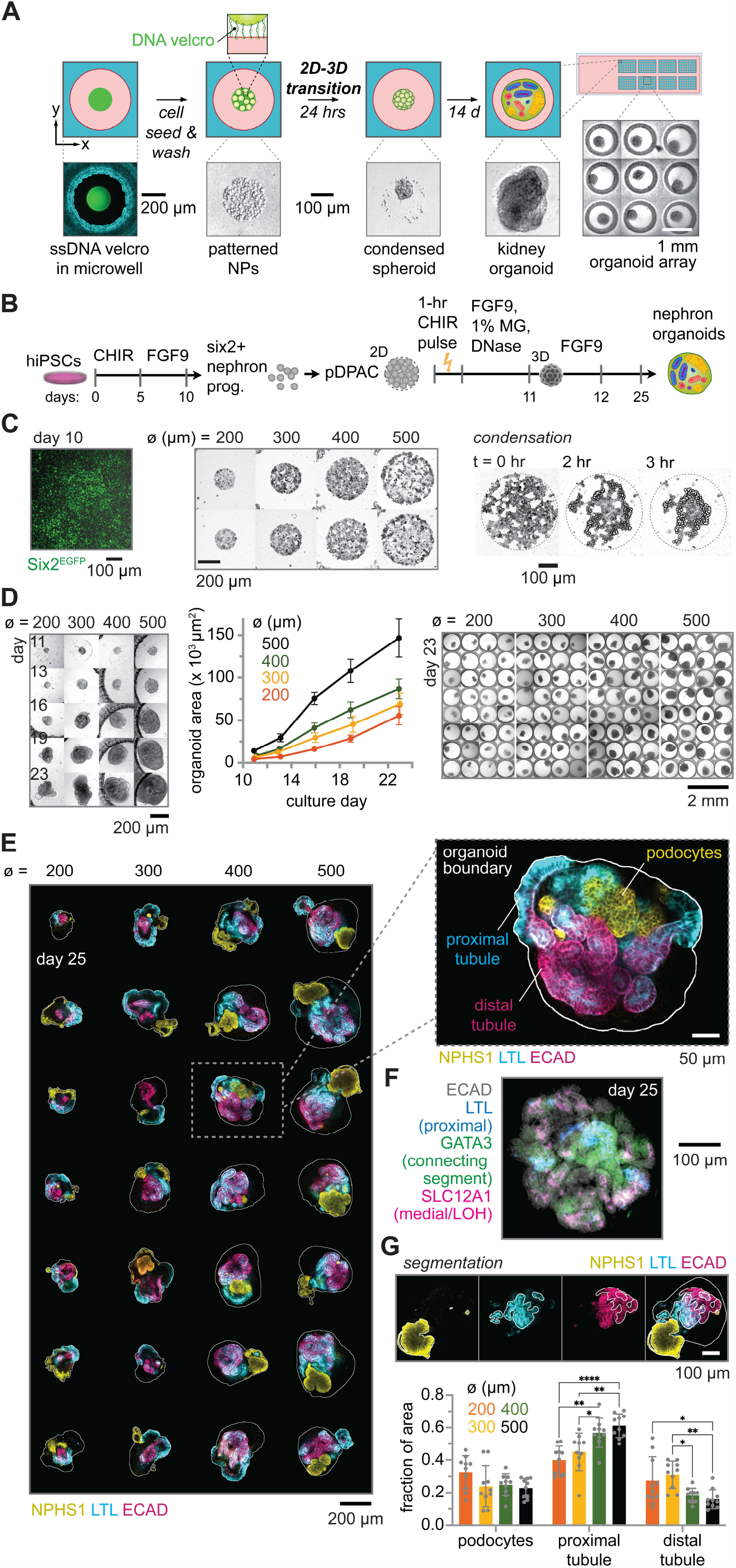
Initial nephron progenitor number biases the emergence of cell types along the early nephron proximal-distal axis. (**A**) Schematic of assay for example of hiPSC-derived SIX2+ NP lineage patterning and differentiation to nephron organoids. (**B**) Patterning and differentiation timeline. (**C**) *Left*, immunofluorescence of Six2^EGFP^ progenitors prior to pDPAC. *Middle*, brightfield examples after cell patterning for ssDNA feature diameters ø. *Right*, frames from **Movie S3** showing 2D to 3D transition. (**D**) *Left*, timepoints of organoids formed from 2D patterns of different ø. *Middle*, representative growth curves (*n* = 10 organoids per group). *Right*, organoid montage at day 23 timepoint. (**E**) Representative confocal immunofluorescence sections at day 25 endpoint. *Inset*, detail of organoid and cell lineages. (**F**) Similar organoid from cells differentiated for 7 rather than 10 days in 2D prior to patterning. LOH: loop of Henle. (**G**) *Top*, segmentation scheme for cell types in day 25 organoids. *Bottom*, plot of organoid composition (ratio of cell type area to total area of all cell types, ± S.D., 3-7 slices per *n* = 10 organoids per ø, Tukey’s multiple comparisons test, *p < 0.0332, **p < 0.0021, ***p < 0.0002, ****p < 0.0001).

These data validated precise 2D cell patterning and translation into 3D spheroids of controlled size and composition that persisted after extended culture.

We next tackled kidney organoid culture. We produced SIX2+ nephron progenitors (NPs)^17,18^ (**Fig. 1C**) lacking mature lineage marker expression prior to patterning (**Fig. S6**). We patterned NPs in microwells using pDPAC, triggered a 2D-to-3D transition by cleaving ssDNA tethers using DNAse, and continued differentiation (**Fig. 1A,B**). Cells successfully condensed into single organoids within ∼4 hrs (**Fig. 1C, Movie S3**) in the presence of either 1% Matrigel or human laminin 521 (**Fig. S7**). 233 of 240 wells (97%) contained single organoids and only 3 (1.3%) were empty at the differentiation endpoint. In contrast, cells passively seeded by gravity typically formed multiple rather than single organoids after 6 days (**Fig. S8**).

To determine if initial cell number could be used to gain control over organoid size and cell composition, we patterned NPs on ssDNA features with diameters ø of 200–500 µm. After 15 days of differentiation, organoid area faithfully reflected differences in initial pattern diameter (**Fig. 1D**). We wondered if the starting pattern size would also change differentiation outcomes. Organoids expressed markers for podocytes (NPHS1/nephrin), proximal tubule (LTL), medial/loop of Henle (SLC12A1), distal tubule (ECAD+ LTL-), and connecting segment (GATA3) nephron cells (**Fig. 1E**) and surrounding stromal-like cells (MEIS 1/2, **Fig. S9**) at the endpoint. Our culture system was compatible with varying protocols, e.g. shortening differentiation from 10 to 7 days before pDPAC and applying a CHIR pulse (**Fig. 1F**)^9^. We segmented day 25 immunofluorescence z-stacks and quantified cross-sectional areas for podocytes, proximal tubule, and distal tubule (**Fig. 1G, Fig. S10**). The starting pattern size had a striking impact on organoid composition. In particular, the representation of proximal tubule as a % of all non-stromal structures significantly increased from 40% ± 8.6% to 61% ± 7.3% from pattern size ø of 200 to 500 µm (± S.D., *n* = 10 organoids per ø). Thus, micropatterning control of initial nephron progenitor number offers significant control over organoid size and markedly affects cell differentiation.

Leveraging pDPAC multiplexing, we next co-patterned NPs and ureteric bud (UB) tip cells since reciprocal signaling between them is crucial to setting kidney size and nephron endowment during kidney morphogenesis^19^. We created mosaic organoids ranging in initial ssDNA area ratios (1:5, 1:2, 2:1, 5:1), directing independent adhesion of hiPSC-derived NPs and UB tip cells to a constant 300-µm square pattern (**Fig. 2A**). UB tip cells were trans-differentiated from distal nephron cells^18^ (**Fig. S11**), forming ruffled organoids with appropriate expression of GATA3, RET (**Fig. 2B**), and additional markers (ECAD, cytokeratin) consistent with UB identity^18^ (**Fig. S12**). Patterned cell ratios differed from nominal ssDNA area ratios by 11.0 % + 5.2% (± S.D., *n ≥* 4 patterns per ratio) (**Fig. S13**). Mosaic NP/UB tip cell patterns created with pDPAC successfully condensed into 3D spheroids within 24-48 hours after DNase treatment (**Fig. 2C**). Next, we scanned culture parameters to find suitable induction/culture conditions for mosaic NP/UB organoids (**SI Note 3**). Interestingly, when NPs were exposed to a 60 min, 7 µM CHIR pulse prior to pDPAC, NP and UB populations sorted, forming ‘core-shell’ morphologies mimicking the *in vivo* interface (**Fig. 2C**). However, without the CHIR pulse, sorting was inverted, with NPs forming cores (**Fig. 2C, *middle*)**. A shift in NP cadherin expression via canonical Wnt signaling could explain the change in cell sorting outcome as dictated by differential adhesion^20^. Our data demonstrate successful integration of distinct hiPSC-derived progenitor lineages into mosaic organoids after cell patterning.

**Fig. 2:**
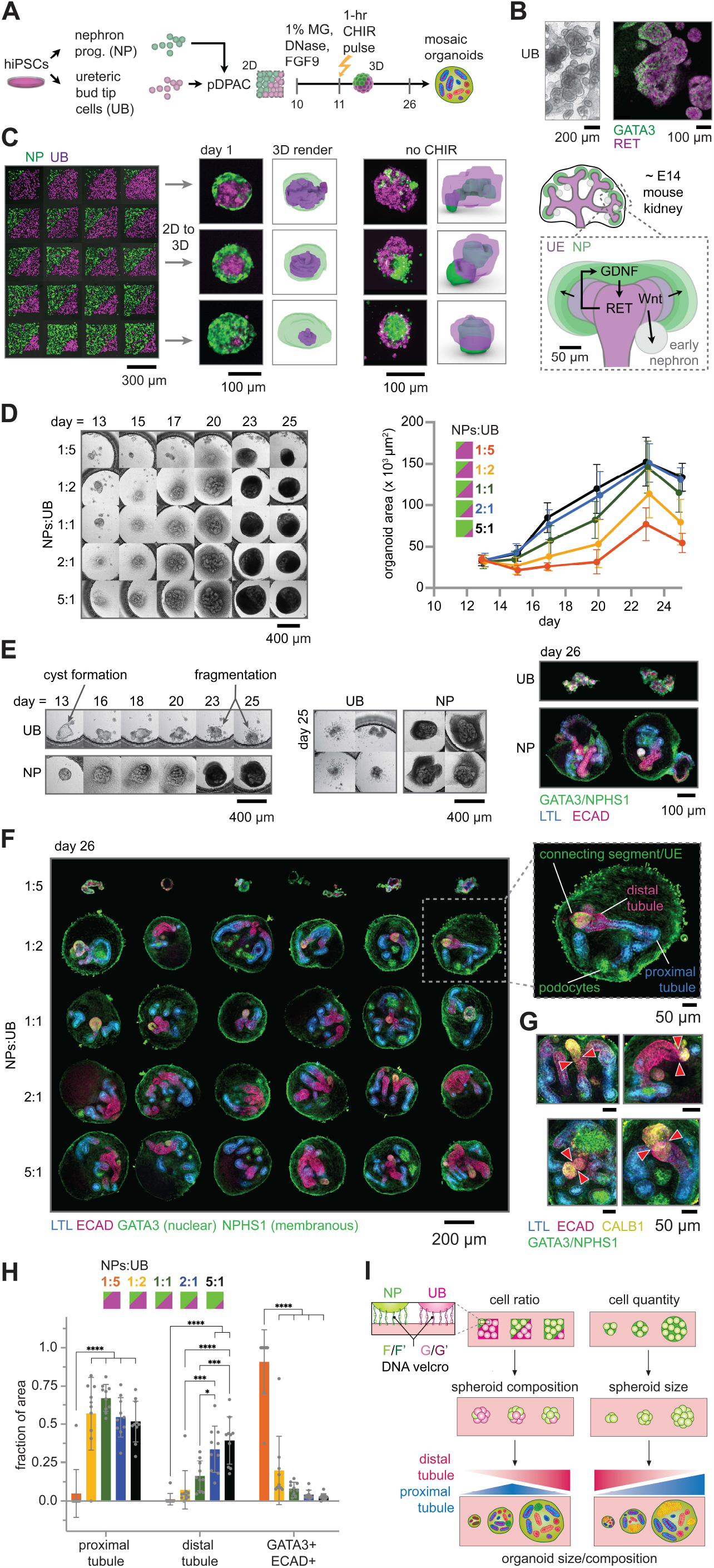
UB tip cell/NP mosaic organoids model the nephrogenic niche interface and trigger cell ratio-dependent emergent patterning. (**A**) Patterning and differentiation timeline. (**B**) *Left*, confocal micrographs of GATA3+ RET+ UB tip cell organoids. (**C**) Fluorescently-labeled UB tip cell and NP co-patterns, and transition to 3D culture +/-NP CHIR-pulse prior to patterning. *Right*, schematic of *in vivo* niche geometry and signaling. (**D**) *Left*, Time-point images of mosaic organoids formed from 2D patterns of different NP:UB cell ratios. *Right*, representative growth curves (*n ≥* 6 organoids per group). (**E**) Example NP-only and UB tip cell-only organoid controls, *Left*, over time, *Middle*, at culture day 25, and *Right*, immunostained as in (**F**). (**F**) Representative mosaic organoids at endpoint (day 26). (**G**) Example Calbindin+ (CALB1+) structures. Arrowheads denote fused (*top*) and unfused (*bottom*) junctions between CALB1+ and distalized nephron structures. (**H**) Mosaic organoid composition (ratio of cell type area to total area of all cell types measured) vs. NP:UB cell pattern area ratios (± S.D., 1-8 slices per *n* = 10 organoids per ratio, Tukey’s multiple comparisons test, *p < 0.0332, **p < 0.0021, ***p < 0.0002, ****p < 0.0001). (**I)** Summary schematic.

We next sought to understand the long-term influence of initial cell ratio on organoid morphology and composition. Mosaic organoid growth increased with increasing NP:UB cell ratio, implying higher proliferation of NPs relative to UB (**Fig. 2D**). Indeed, NP-only control organoids grew steadily compared to UB-only controls, which in the absence of NPs, formed cystic structures that inflated, deflated, and fragmented (**Fig. 2E**). Substrate-adherent, stromal-like cells were presumably derived from the NP cell population, as they were not found in UB-only controls (**Fig. 2E**). We performed confocal immunofluorescence analysis of day 26 mosaic organoids using markers of connecting segment/UE (GATA3+ ECAD+), Calbindin1+ tissue (CALB1+), distal tubule (ECAD+ LTL-GATA3-), proximal tubule (LTL+), and podocytes (NPHS1+) (**Fig. 2F**). Through manual image segmentation, we found that only 1 of 10 organoids derived from an initial NP:UB tip cell patterning ratio of 1:5 produced NP-derived proximal tubule, distal tubule, or podocytes. Moving to a 1:2 initial NP:UB tip cell ratio rescued nephron structure formation in 9 of 10 organoids, suggesting that a minimum number or ratio of NPs is required for nephrogenesis. The proportion of proximal tubule structures increased and then decreased with increasing initial NP cell ratio (**Fig. 2H, Fig. S14A**), peaking at 1:1. This suggests a ‘goldilocks’ NP:UB cell ratio that maximizes proximal tubule, indicating an interaction between cell autonomous and non-autonomous cues from the UB in its induction^9,21^. Distal tubule monotonically increased with increasing initial NP:UB ratio, while GATA3+ ECAD+ connecting segment/ureteric epithelium structures decreased (**Fig. 2H, S14A**). Interestingly, we saw no significant differences in the proportion of CALB1+ structures and only minor differences in representation of podocytes across all starting ratios (**Fig. S14B**). While not specific to UB epithelium *in vivo*, CALB1 is highly upregulated in it^18^. In rare instances (3/50 organoids), we observed complete *in vivo*-like fusion of CALB1+ structures with GATA3+ ECAD+ connecting segment/UE, along with proper distal-to-proximal segmentation, and an equivalent number of CALB1+ structures juxtaposed but not fused to GATA3+ ECAD+ structures (**Fig. 2G**). In controls, CALB1+ structures were not observed, implying that NP and UB interaction is required for CALB1 upregulation. These data show that initial NP:UB cell ratio in mosaic kidney organoids modulates compositional outcomes, shifting the representation of tissues along the proximal-distal axis while amplifying emergent inductive and connectivity phenomena.

Here we advance precision cell patterning for 3D organoid culture to improve throughput, imaging accessibility, and size homogeneity, while offering new capabilities for control over morphogenetic outcome (**Fig. 2I**). This expands upon cell aggregation by centrifugation, agitation, or seeding in microwells^1,3,11,12,22^, which lack precise control over organoid size and composition, and cell patterning in 2D that lacks a transition to 3D self-organization^23^. We contribute an integrated ssDNA-based cell patterning and long-term microwell culture platform that is compatible with hiPSC-derived cell lineages. Our technology offers opportunities for automation and tracking that enable studies of growth, cell sorting, segmentation, and fusion of different structures/cell populations. These advances enable controlled initial conditions and downstream screens applicable to diverse organoid and synthetic embryo systems.

## Acknowledgements

We thank Louis Prahl for advice on DNA patterning conditions, Ananya Gupta and Wenli Yang for advice on hiPSC maintenance, and Sarah Howden and Melissa Little for advice on hiPSC differentiation protocols. This research was conducted in part at the Singh Center for Nanotechnology at the University of Pennsylvania, which is supported by the NSF National Nanotechnology Coordinated Infrastructure Program under grant NNCI-2025608. This work was supported by an NSF GRFP award (CMP), NIH NIGMS MIRA R35GM133380 (AJH), and an NSF CAREER award 2047271 (AJH) and was partially supported through the University of Pennsylvania Materials Research Science and Engineering Center (NSF MRSEC, DMR-2309043).

## Methods

### Cell lines

Madin-Darby canine kidney cells (MDCKs) expressing H2B-VFP or H2B-iRFP were maintained as previously described^16^. Briefly, cells were passaged using 0.25% Trypsin (Thermo Fisher Scientific, 25300056) and cultured at 37°C and 5% CO2 in T-175 flasks (Corning) in minimum essential medium (MEM, with Earle’s Salts and L-glutamine, Corning 10-010-CM) and 10% fetal bovine serum (Corning 35-010-CV).

Nephron progenitor (NP) and ureteric bud (UB) tip cells were derived from human hiPSCs similarly to previous reports^12,18,24^. Six2^EGFP^ reporter hiPSCs^17^ were maintained at 37°C and 5% CO2 on plates coated with Matrigel (hESC-qualified, Corning, 354277) in mTeSR+ (STEMCELL Technologies, 100-0276) and passaged every 3-4 days using Gentle Cell Dissociation Reagent (STEMCELL Technologies, 100-0485) for clump passaging, or Accutase (Thermo Fisher Scientific, A1110501) for single-cell passaging. Genomic integrity was confirmed by molecular karyotyping through the Induced Pluripotent Stem Cell Core, Penn Institute for Regenerative Medicine.

### hiPSC-derived NPs

hiPSCs were lifted and dissociated to single-cell suspension with Accutase at 37°C, counted with a hemocytometer and seeded at 65,000 cells per well in a 6-well culture plate coated with Laminin 521 (BioLamina, 77003). The following day, induction of intermediate mesoderm began using 7 µM CHIR99021 (R&D Systems, 4423), a Wnt agonist, in TeSR-E6 medium (Stem Cell Technologies, 05946) for 5 days. Media was then swapped to 200 ng ml^-1^ FGF9 (R&D, 273-F9-025) and 1 µg ml^-1^ heparin (Sigma Aldrich, H4784) in TeSR-E6 for 5 days to induce nephron progenitors. On day 10, cells were dissociated with Accutase at 37°C to single-cell suspension, diluted 5x with TeSR-E6, and pelleted at 200 g for 3 min.

### hiPSC-derived UB tip cells

UB tip cells were trans-differentiated from NP cultures derived similarly to the above, except with two adjustments thought to favor anterior intermediate mesoderm from which the ureteric epithelium derives *in vivo* and/or more distal nephron cell identity^18^. Specifically, the CHIR step was reduced to 3 days, and 200 ng ml^-1^ FGF9 was replaced with 600 ng ml^-1^ FGF2 for 4 days.

Cells were then dissociated with TrypLE at 37°C to single-cell suspension, diluted 5x with TeSR-E6 and pelleted at 300 g for 2.5 min. Media was aspirated and cells resuspended as a dense slurry in residual media. 2 µl of this slurry was spotted onto 0.4 µm polyester transwell membranes in 6-well plates with 4 organoids per membrane. Organoids received a CHIR pulse by culturing at the air-liquid interface for 1 hr in the presence of 7 µM CHIR99021 in TeSR-E6 medium in the lower transwell compartment (1.2 ml well^-1^). Media was swapped to TeSR-E6 supplemented with 600 ng ml^-1^ FGF2 and 1 µg ml^-1^ heparin for 5 days, and then to TeSR-E6 supplemented with 0.1 µM TTNPB (a retinoic acid analogue, Tocris, 0761) for 13 days.

To begin transdifferentiation, day 20-25 organoids were dissociated with 200 µl 1:1 TrypLE (Thermo Fisher Scientific, A1285901):Accutase per organoid, occasionally agitating by gentle vortexing and trituration. The suspension was then diluted 5x with TeSR-E6 medium, pelleted for 3 min at 300 g, and resuspended at 6 x 10^6^ cells ml^-1^ in 200 ng/ml FGF2, 3 μM CHIR99021, 0.1 μM TTNPB, 10 μM Y-27632 and 100 ng ml^-1^ GDNF (‘UE medium’). At least 6 x 10^5^ cells (in 100 µl media) were then plated per well of a 6-well polyester Transwell plate. 1.2 ml of 1:1 growth factor-reduced Matrigel:UE medium were added on top of each transwell, and 2.3-2.5 ml of UE medium were added to each basolateral compartment. For passaging, Matrigel was first digested using 3 ml of dispase per well (PluriSTEM™ Dispase-II, 1 mg/mL, Millipore Sigma, SCM133) and incubated for 10 min in a 15 ml conical tube at 37°C, vortexing gently at 5-min intervals. 9 ml of TeSR-E6 medium was then added and the mixture was pelleted at 500 g for 5 min. Partially degraded matrigel was then aspirated off and the pellet subjected to continued dissociation with 2 ml of 1:1 TrypLE:accutase at 37°C for 15-25 min with periodic gentle vortexing. 10 ml of DPBS with 2% FBS was added and cells pelleted at 500 g for 3 min, followed by resuspension as before and Transwell plating. At least one passage was required to obtain a 99% pure UE population, since UE propagation was favored over off-target cells in UE medium.

Freezing medium for cell storage consisted of 10% DMSO, 40% knockout serum replacement (Thermo Fisher Scientific, 10828010) 50% TesR-E6, and 10 µM Y-27632.

### Fabrication of photoactivatable substrates for single-stranded DNA (ssDNA) patterning

Photoactivatable polyacrylamide (PPA) gels were fabricated on glass slides as previously described^16^. Briefly, according to manufacturer guidelines, SU-8 2025 photoresist (MicroChem, Y111069) was coated at a thickness of 30 μm onto mechanical-grade silicon wafers (University Wafer) using a digital spin coater (INSTRAS Scientific, SCK-300P). The wafer was soft baked on a hotplate (Fisher, SP88857200) at 65°C for 1 min and 95°C for 4 min. After cooling, a Mylar mask, printed with a double-rail pattern at 20,000 d.p.i. (CAD/Art Services), was laid onto SU-8-coated wafers and exposed for 30s to 365-nm UV light from a mounted LED (ThorLabs M365LP1, ACL7560U, SM3V10) at 10 mW/cm^2^, as measured by a light meter (Thorlabs PM100D, S120VC). The Mylar mask was designed to pattern photoresist rails that would run the 75-mm edges of standard glass slides, creating a 30-µm gap between the slide and the silicon wafer for molding of a PA gel layer. Wafers were baked at 65°C for 1 min and 95°C for 3 min after exposure, allowed to cool to RT, developed in an SU-8 developer solution (MicroChem, Y020100), and washed with isopropanol, followed by acetone. Two milliliters of hydrophobic dichlorodimethylsilane (DCDMS, Sigma-Aldrich, 440272) were deposited on wafers *in vacuo* for 15 min. Silanized wafers were washed thoroughly with deionized (DI) water and dried under a compressed airstream. Between PA gel fabrications, silicon wafers were washed with 0.1% Triton in DI water, followed by DI water.

Plain 75 x 25 mm glass microscope slides (Corning, 2947-75X25) were rinsed in 0.1% Triton in DI water to remove surface grease and dried under an airstream. Slides were silanized with 3-(Trimethoxysilyl)propyl methacrylate (Sigma-Aldrich, 440159-100ML) to create a monolayer of methacrylate functional groups according to established protocols^25^. Silanized slides were placed, functionalized-side-down, on patterned silicon wafers and manually aligned with their 75-mm edges against SU-8 rails. PA gel precursor solutions were made out of the following: 7% T (w/v% total acrylamides), made from a 30% T, 2.7% C (w/w% of the cross-linker *N,N-*methylenebisacrylamide) stock (Sigma-Aldrich, A3699); 3 mM benzophenone-methacrylamide (N-[3-[(4-benzoylphenyl) formamido]propyl] methacrylamide, BPMAC, PharmAgra) from a 100 mM stock in DMSO, 0.06% SDS (Bio-Rad, 161-0301), 0.06% Triton X-100 (Fisher, BP151), 0.06% ammonium persulfate (APS, A3678, Sigma-Aldrich), 0.06% tetramethylethylenediamine (TEMED, T9281, Sigma-Aldrich), 1x DPBS (Ca^2+^/Mg^2+^-free, Thermo Fisher Scientific, 14-200-075), and DI water. Partial precursors were made of acrylamides and DPBS and degassed *in vacuo* in an ultrasonic bath (15-337-411, Fisher) for 1 min.

Detergents SDS and Triton were added, followed by BPMAC, and finally, APS and TEMED catalysts. Using a standard 200-μl pipette, ∼150 μl of precursor solution were then injected into the gap between the methacrylate-functionalized glass slide and silicon wafer. Precursor spread through the gap for ∼30 s, and the slide was slid along the length of the rails to allow any remaining bubbles to escape through its 25-mm ends. Excess precursor was removed using a Kimwipe, ensuring flush contact of the slide with the SU-8 rails. The slide was then left alone for 25 min to allow for additional PA polymerization.

Following polymerization, 2 ml of DPBS were pipetted against one 25-mm slide edge; at this side, the slide was carefully levered from wafer using a razor blade, with DPBS wicking beneath the gel to aid release. Fabricated slides were stored in DPBS at 4°C for up to 1 week before ssDNA was photopatterned.

### Design and fabrication of photomasks for ssDNA patterning

CAD files for ssDNA spot and square array patterns were designed in LayoutEditor, then finalized and converted to Heidelberg format files in BEAMER software. Custom 5”x5” photomasks were fabricated in a cleanroom facility: Spot/square arrays were exposed on blank chrome-on-quartz masks using a DWL 66+ laser lithography system (Heidelberg Instruments) to direct write on 0.5-µm thick coatings of IP3500 positive photoresist. The masks were developed in CD-26 solvent and chromium etched. Then, remaining resist was stripped using Dow MICROPOSIT™ Remover 1165. Masks were rinsed with acetone and IPA and air-dried.

### Patterning of ssDNA on PA gels for pDPAC

Oligos (IDT, 5’-T_20_-X_20_-3’) were patterned on photoactivatable PA gels attached to glass slides. Two oligos were used: “F,” where X20 is 5’-AGAAGAAGAACGAAGAAGAA-3’ and “G,” where X_20_ is 5’-AGCCAGAGAGAGAGAGAGAG-3’. In a glove box (Bel-Art, H50028-2001) filled with an atmosphere of medical-grade nitrogen, PPA gels were dried under a nitrogen stream. For each slide, a 400-µl solution of 0.25 mM (for MDCK patterning) or 0.375 mM (for hiPSC-derived cells) oligo in 1x DPBS was degassed, moved into the glove box, and sparged with a nitrogen stream for ∼30s. The oligo solution was pipetted onto the patterned chrome side of a photomask, and the slide was laid—gel-side-down—onto the liquid bead of oligo solution; starting at an ∼45-degree angle, the slide was carefully lowered from one edge to the other, allowing the oligo solution to wick across the PPA gel without introducing bubbles. After manually aligning the slide above the mask pattern and letting it rest for ∼1 min, excess oligo solution was removed with a Kimwipe, immobilizing the slide to the mask. The slide-mask sandwich was removed from the glove box, flipped, and exposed to 254-nm light in a UV oven (Spectrolinker XL-1000 UV Crosslinker, Spectronics Corporation) for 2 min at ∼9 mW/cm^2^. After 254-nm UV light exposure, 2 mL of 0.1% SDS in DI water were pipetted against one 25-mm slide edge, and the slide was carefully levered from the mask using a razor blade. To remove unadhered oligo, the slide was then soaked for 10 hrs in 15 mL of 0.1% SDS in DPBS in a 15-cm petri dish. Fresh solution was used to rinse once more for 20 min, followed by two 20-min washes in DPBS only to remove SDS. In the case of patterning a second oligo, UV exposure to pattern the first oligo was reduced to 110 s, and the slide was dried under an airstream following wash steps. The alignment fiducial marks of the first oligo (G) were stained with a 0.2 μM solution of a custom, fluorescently-tagged complementary oligo (5’-/56-FAM/CTCTCTCTCTCTCTCTGGCT-3’, IDT) in DPBS for 10 min and rinsed in a petri dish of DPBS for 5 min prior to drying and application of the second oligo (F) in nitrogen. Upon removal of the slide-mask sandwich from the glove box, the slide was manually aligned on the second mask pattern to the stained oligo G fiducial marks under a 470-nm blue light using an inverted microscope (Eclipse Ts2-FL, Nikon) (**Fig. S5B**). Following 2-min UV light exposure to pattern F oligo, the slide was again carefully levered from the mask and free F oligo was removed through the previously described washing steps. The slide was stored in fresh DPBS at 4°C and dried under an airstream prior to the attachment of polydimethylsiloxane (PDMS) microwells.

### PDMS microwell fabrication

PDMS sheets patterned with through-holes were molded using 3D-printed pillar arrays. Conical frustum pillar arrays were designed in SolidWorks such that their positions coincided with the layout of the 8 chambers of cell culture slides (MatTek Corporation, CCS-8) as well as ssDNA patterns. The arrays were then printed in grey resin (Formlabs, RS-F2-GPGR-04) by a 3D printer (Formlabs Form 3), at a printing resolution of 25 μm. Pillar arrays were post-processed by rinsing in 100% isopropanol (Form Wash instrument, Formlabs), removing from supports, and drying for at least 1 hr. To reduce bowing of the mold, curing processes of UV exposure and baking were done separately. The mold was exposed to UV in a Form Cure instrument (Formlabs) without heat. It was then placed under a glass slide and 500-g weight (Troemner, 61055S) and baked for 24 hrs at 60°C. To remove any residual uncured resin in the mold, which could inhibit PDMS curing, it was soaked in isopropanol for 15+ hrs and dried.

PDMS sheets were then molded against 3D printed pillar arrays. A 10:1 base to catalyst solution of PDMS silicone rubber (Sylgard 184, Ellsworth Adhesives, 2065622) prepolymer was thoroughly mixed and degassed in a vacuum chamber. Approximately 3 ml of PDMS prepolymer were poured onto the pillar array. A metal spatula was used to spread and level the prepolymer. The tops of the pillars were blown with a gentle airstream. PDMS was baked at 40°C (Heratherm IMH100 Advanced Microbiological Incubator, Thermo Fisher Scientific, 51028067) for 48 hrs. Following curing, the tops of pillar arrays were firmly rubbed to remove any residual PDMS; a microporous cosmetic sponge was soaked in isopropanol and wrung out, then used to wipe the tops of the pillars. PDMS sheets were then demolded and washed in 100% isopropanol for 24 hrs and air-dried. Before reuse, pillar arrays were rinsed with 100% isopropanol.

### PDMS microwell passivation

Similar to published methods^26–29^, PA was grafted onto PDMS through-hole sheets to passivate against nonspecific cell and protein adhesion during culture. Dry PDMS sheets were placed on a glass microscope slide. Each sheet was then plasma-treated with a hand-held high frequency generator (Electro-Technic Products, Inc., Model BD 10A) in a raster motion for 30 seconds on each side and submerged for 15 min in a 10% v/v solution of 3-(Trimethoxysilyl)propyl methacrylate in acetone. Sheets were then soaked in a 5% w/v solution of benzophenone (Sigma-Aldrich, B9300-25G-A) in acetone for 15 min. In a nitrogen atmosphere, PDMS sheets were flipped on glass slides to remove excess solution and thoroughly dried under nitrogen. They were placed on a slide, top-side (larger through-hole diameter) up. Approximately 1.5 mL of a degassed and nitrogen-sparged solution of 15% w/v acrylamide monomer (Fisher Scientific, BP170-500) in DI water was pipetted onto PDMS sheets. A quartz slide (Thermo Fisher Scientific, AA42297KG) was laid on the PDMS sheets. The quartz-PDMS-glass sandwich was exposed to 254-nm light in a UV oven for 10 min. PDMS sheets were then washed alternately in 70% ethanol, DI water, and 70% ethanol again for 30 min each, and air-dried.

### PDMS microwell alignment and adhesion to PA gels

The DNA-patterned PA gel was dried under an airstream. The four corners and center of each chamber pattern were stained with a solution of 20x SYBR Gold (Invitrogen, S11494) in DI water for 10 min. To remove non-adhered SYBR, the slide was soaked in a petri dish of DPBS for 10 min and air-dried.

For alignment and attachment of PDMS sheets to the PA base gel, the side of each PDMS sheet, which was not cast directly against the 3D-printed mold (i.e., smaller through-hole diameter side), was plasma-treated for 1.5 min in a raster pattern with a hand-held high frequency generator. The PDMS sheet was immediately submerged in DI water. The PDMS sheet was transferred, plasma-treated side down, onto the PA gel in the approximate region of a chamber’s array of ssDNA patterns. SYBR-stained DNA patterns were illuminated using collimated, 470-nm blue light (Thorlabs, COP1-A and M470L4) mounted on a ring stand and visualized through a stereo microscope (Nikon, SMZ800N).

Before the water dried, a stainless-steel probe (Fine Science Tools, 10140-04) was used to manually align each PDMS through-hole sheet, such that DNA patterns were centered in each microwell. The PDMS sheets immobilized upon complete evaporation of DI water. Once each PDMS sheet had been aligned and adhered, a small piece of aluminum foil was laid on the PPA/PDMS microwell slide, followed by a large glass slide (Corning, 2947-75X50) and a 500-g weight (Troemner, 61055S). The slide was then baked at 70°C for 16 – 18 hrs to anneal the PDMS to the PA gel.

### Cell patterning in microwells

MDCKs and hiPSC-derived NPs and UE cells were patterned on ssDNA features within fabricated PDMS/PPA composite microwells. Prior to cell seeding, each microwell slide was soaked in 3% bovine serum albumin in DPBS for 1 hr, rinsed with two changes of DPBS in a petri dish, and stored in fresh DPBS at 4°C until it was needed for cell seeding. Directly prior to seeding, the gel was soaked in 70% ethanol for 30 min and rinsed with two changes of sterile DPBS.

In some experiments, cells were labeled with CellTracker dyes prior to lifting for cell patterning. Lyophilized CellTracker Red (Thermo Fisher Scientific, C34552), Deep Red (Thermo Fisher Scientific, C34565), and Green CMFDA (Thermo Fisher Scientific, C7025), were each resuspended in DMSO to the manufacturer’s recommended concentrations. Each was then diluted to 1 µM in serum-free MEM. Adherent cells were incubated in CellTracker medium for 30 min at 37°C. CellTracker medium was then removed and the cells were washed with DPBS.

For MDCK patterning, cells grown to ∼80% confluency in T-175 polystyrene culture flasks were washed with DPBS and incubated at 37°C in 0.25% Trypsin-EDTA for ∼10 min to lift them. MDCKs were resuspended in culture media and centrifuged at 200 g for 3 min at 4°C. They were then washed twice by resuspending in 10 mL of DPBS and re-pelleted by centrifugation. MDCKs were resuspended in 100 µL of DPBS in 1.5-mL Eppendorf tubes (1 tube per T-175) with 1 mM EDTA (Thermo Fisher Scientific, 15575-038) and labeled with lipid-DNAs (custom syntheses, OligoFactory, Holliston, MA): “universal anchor” (5’-TGGAATTCTCGGGTGCCAAGGGTAACGATCCAGCTGTCACT-C24 lignoceric acid-3’), a lipid-conjugated ssDNA, was added to each Eppendorf tube from a 100 µM stock in DI water to a final concentration of 2.5 µM. Then, lipid-conjugated ssDNA “universal co-anchor” (5’-C16 Palmitic acid-AGTGACAGCTGGATCGTTAC-3’) was added to a final concentration of 2.5 µM, followed by 2.5 µM final concentration of “adhesion strand” DNA (5’-CCTTGGCACCCGAGAATTCCA-T19-Y20-3’, where Y20 is the reverse complement of the X20 sequence patterned on the pDPAC slide)^16,30,31^. Each oligo was added in succession to the 100-µL reaction; an 8-min incubation step under gentle agitation on a vortex set at very low speed (∼5 Hz) followed each addition. After adding the series of 3 oligos, cells were washed 3 times in 1 mL of DPBS with 1 mM EDTA by pelleting through centrifugation and aspirating off DPBS. At the end of labeling and washing, 600 µL of DPBS with 1 mM EDTA were added to each Eppendorf tube, and cells labeled with the same ssDNAs were combined and placed on ice.

Excess DPBS was poured off each ssDNA-patterned slide. Using a 200-μl pipette, cell suspension was added dropwise over the microwells, such that it fully covered all DNA patterns. Slides rested in a petri dish on ice for 5 min as cells settled in microwells. Then, each slide was dipped repeatedly into a cold bath of DPBS with 1 mM EDTA to remove unpatterned cells. Cell patterns were intermittently checked on an inverted microscope between washes until unpatterned cells had been fully removed from the microwells. For dual MDCK patterns, the second oligo-labeled cells were then added dropwise to the slide and settled for 5 min, and the wash steps to remove nonspecific cells were repeated.

For hiPSC cell-derived pDPAC, some changes were made to the patterning protocol. First, cells were maintained at RT in TeSR E6 medium with 100 µM Y-27632 (Tocris, 1254) throughout the oligo functionalization steps. 2.5 x 10^7^ NPs were functionalized with 5 µM each of universal anchor, co-anchor, and adhesion strand F’. Due to the larger surface area of each UB tip cell, 1.7 x 10^6^ UB tip cells were functionalized with 5 µM each of universal anchor, co-anchor, and adhesion strand G’. Following, the 3 cell pellet washes to remove excess oligo as well as cell patterning and postpatterning microwell washes were carried out in RT DPBS. Each oligo reaction of cells was resuspended in 500 ul of DPBS before patterning. For dual NP/UE pDPAC, UE cells remained in the last oligo addition and were only washed 3 times and resuspeded once NPs had been patterned and the microwells thoroughly washed of unadhered cells. UE was then patterned second, followed by microwell washing.

### Device assembly and culture

After pDPAC, slides were loaded into bases of 8-well cell culture chamber slides (MatTek, CCS-8). Gaskets were removed from the manufacturer’s provided glass slides and inserted into the grooves of the polystyrene chambers. The chambers were then aligned over the microwell slides and the chamber bases were clamped in place.

For NP only organoids, patterned cultures were incubated in a pulse of TeSR-E6 with 7 µM CHIR99021 and 10 µM Y-27632 for 1 hr and exchanged to TeSR-E6 with 10 µM Y-27632, 1% Matrigel, 200 ng ml^-1^ FGF9, and 1 µg ml^-1^ heparin for ∼ 3 hr until cells spread on ssDNA patterns and formed visible cell contacts. 15 µl of TURBO DNase (Fisher Scientific, AM2238) were added to each chamber to cleave ssDNA and initiate aggregate formation. The following day, medium was swapped to TeSR-E6 with 200 ng ml^-1^ FGF9 and 1 µg ml^-1^ heparin. Chamber slides were placed on an orbital shaker at 60 rpm for the rest of the culture period (14 days). The next day, medium was swapped to TeSR-E6 only and exchanged every 2 days for 13 days.

For UE/NP mosaic organoids, patterned cells were incubated in TeSR-E6 with 10 µM Y-27632, 1% Matrigel, 300 ng ml^-1^ FGF9, 0.5 µg ml^-1^ heparin and 2% FBS. After ∼ 3 hrs, cells had formed visible cell junctions, at which time 15 µl of TURBO DNase were added to each chamber to initiate transition to 3D culture. Twenty-four hrs later, cultures received a pulse of TeSR-E6 with 7 µM CHIR99021 and 10 µM Y-27632 for 1 hr at 37°C. Medium was then swapped to TeSR-E6 with 300 ng ml^-1^ FGF9 and 0.5 µg ml^-1^ heparin for 24 hrs. Organoids were then maintained in plain TeSR-E6 on an orbital shaker at 60 rpm for the rest of culture, with medium exchanged every 2 days for 13 days.

### Immunofluorescence

Immunofluorescence staining and imaging was performed as previously described^32^, using protocols adapted from Combes *et al*. and O’Brien *et al*.^*33,34*^. Briefly, 15 days post pDPAC, organoids were fixed in 4% paraformaldehyde in DPBS for 45 min, washed three times for 5 min per wash in DPBS, and blocked for 2 hrs at room temperature in PBSTX (DPBS + 0.1% Triton X-100) containing 5% donkey serum (D9663, Sigma). Following, fixed and blocked organoids were incubated in primary and then secondary antibodies in blocking buffer for at least 24 hrs each at 4°C, alternating with 3 washes in PBSTX, with a 30-min wait after the first two PBSTX additions, and a 12 to 24-hr wait after the last PBSTX wash.

Primary antibodies and dilutions included biotin anti-human LTL (1:300, B-1325, Vector Laboratories, RRID:AB_2336558), goat anti-human GATA3 (1:20, AF2605, R&D Systems, RRID:AB_2108571), rabbit anti-human GATA3 (1:500, 5852, Cell Signaling Technology, RRID:AB_10835690), mouse anti-human E-cadherin (1:300, 610181, Biosciences, RRID:AB_397580), rabbit anti-human E-cadherin (1:300, 3195, Cell Signaling Technology, RRID:AB_2291471), rabbit anti-human SLC12A1 (1:300, ab171747, Abcam, RRID:AB_2802126), sheep anti-human Nephrin (1:40, AF4269-SP, R&D Systems, RRID:AB_2154851), goat anti-human RET (1:50, AF1485, R&D Systems, RRID:AB_354820), rabbit anti-human RET (1:200, 3223, Cell Signaling Technology, RRID:AB_2238465), mouse anti-calbindin D-28K (1:500, clone CB-955, C9849, Sigma, RRID: AB_476894), mouse anti-pan-cytokeratin (1:200, clone 11, C2931, Sigma, RRID:AB_258824), and mouse anti-MEIS 1/2/3 antibody (1:200, clone 9.2.7, 39796, Active Motif, RRID:AB_2750570). Secondary antibodies (raised in donkey) were used at 1:200 dilution and included anti-rabbit AlexaFluor 488 (A21206, ThermoFisher, RRID: AB_2535792), anti-mouse AlexaFluor 555 (A31572, Thermo Fisher, RRID: AB_162543), anti-rat AlexaFluor Plus 555 (A48270, Thermo Fisher, RRID: AB_2896336), anti-goat AlexaFluor Plus 647 (A32849, Thermo Fisher, RRID: AB_2762840), and anti-sheep AlexaFluor 647 (A-21448, Thermo Fisher, RRID: AB_2535865).

Finally, DyLight 405-Streptavidin (016-470-084, Jackson ImmunoResearch) was used to stain biotinylated LTL.

### Imaging

Imaging was performed using a Nikon Ti2-E microscope equipped with a CSU-W1 spinning disk (Yokogawa), a white light LED, laser illumination (100 mW 405, 488, and 561 nm lasers and a 75 mW 640 nm laser), a Prime 95B back-illuminated sCMOS camera (Photometrics), motorized stage, 4x/0.2 NA, 10x/0.25 NA and 20x/0.5 NA lenses (Nikon), and a stagetop environmental enclosure (OkoLabs).

### Image analysis

For longitudinal analyses, we selected organoids derived from progenitor patterns that displayed high initial ssDNA patterning fidelity and coverage and low nonspecific background cell adhesion. Immunofluorescence marker quantification was performed from 3-7 z-slices per NP-only organoid and 1-8 z-slices per mosaic NP/UB tip cell organoid recovered from confocal fluorescence micrograph stacks, consisting of the approximate mid-plane, and respective planes at -25 and +25 µm in *z*, with additional 25-µm increments in *z* to span the organoid volume. For each slice, regions attributed to each marker category–podocytes (NPHS1, in NP-only organoids), proximal tubule (LTL), and distal tubule (ECAD+ LTL-GATA3-)–were manually segmented in FIJI. In NP/UB tip cell mosaic organoids, UE/connecting segment (ECAD+ GATA3+), calbindin+ structures, and podocytes made up a small overall proportion of the organoid compared to the distal and proximal tubule and tended to form more spherical compartments that spanned fewer slices in z. Thus, for these tissues, the projected area of each discrete compartment was segmented and measured. We defined the remaining area of the organoid as a stromal-like population, which was supported by positive MEIS1/2 immunostaining.

We then calculated the area fraction of each tissue as the total area of each tissue divided by the total area of all measured, non-stromal-like tissues.

Organoid growth was tracked using brightfield confocal images taken at 2-3-day intervals throughout organoid culture in microwells. All organoids selected for growth analysis had lifted from PA substrates and rounded by their culture endpoints. At each analyzed time point, the maximum projected area of each organoid was manually segmented and measured. In cases of fragmented organoids or organoids that failed to condense to a single organoid per microwell, the projected areas of the total tissue per microwell was summed. For substrate-adhered organoids, areas of apparent epithelialized structures that stood out from surrounding flattened cells were manually segmented and measured.

3D renderings of CellTracker-stained mosaic organoids were generated by manual segmentation of z slices from confocal fluorescence stacks to create binary stacks, exporting as .stl surface objects from FIJI using the 3D Viewer plugin, followed by importing and rendering in Rhino 7 3D modeling software (Robert McNeel & Associates).

## Statistical analysis

One-way analysis of variance (ANOVA) with correction for multiple comparisons using Tukey’s honestly significant difference test was performed in Prism 10 software (GraphPad). Trend analyses were conducted through curve fitting using linear and nonlinear least squares regression methods, also performed in Prism 10 software.

## Notes

### Competing Interest Statement

The authors have declared no competing interest.

