## Supplementary Information for "Highly-parallel production of designer organoids by mosaic patterning of progenitors"

### Supplementary Figures

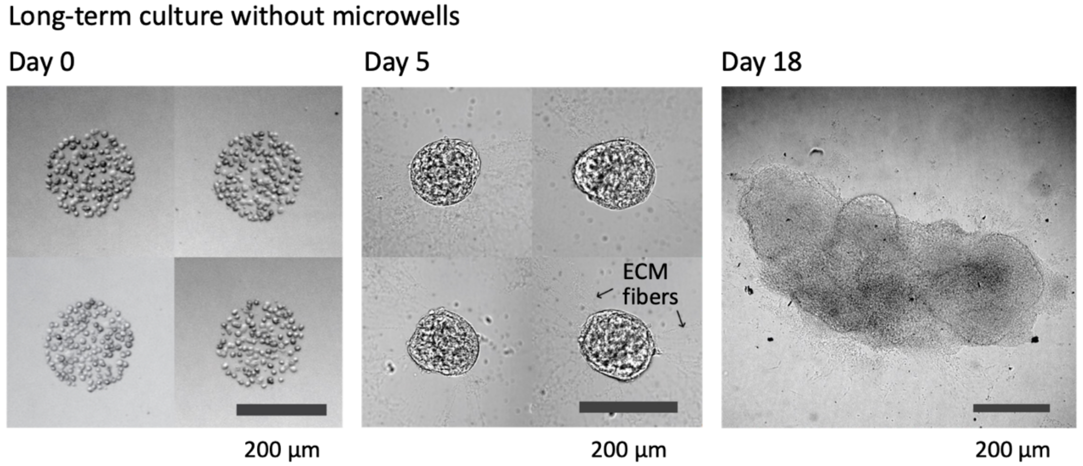


**Fig. S1: Microwells prevent organoid fusion upon extended culture.** Example brightfield micrographs over a differentiation time-course for nephron progenitor organoids patterned in arrays without microwell walls. Organoids appear to interact with ECM fibers, drawing neighboring organoids together into large masses over time.

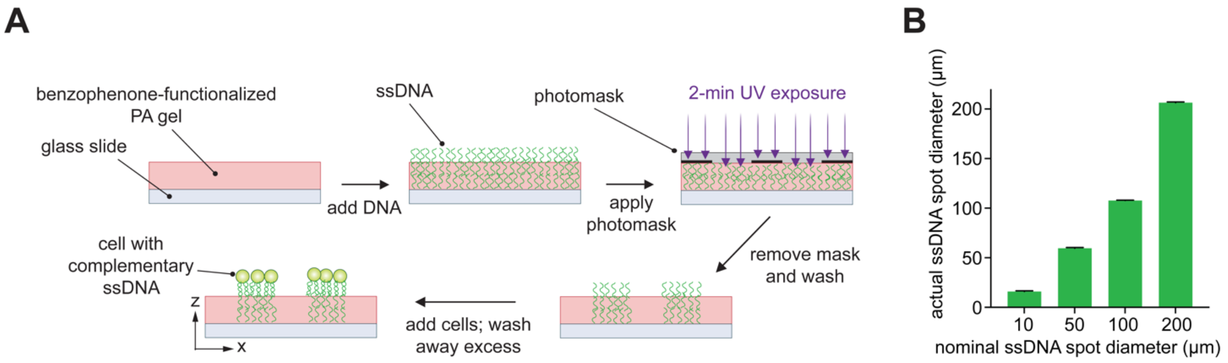


**Fig. S2: ssDNA photopatterning is spatially accurate and precise.** (**A**) Schematic of pDPAC: a solution of ssDNA is introduced to a photoactive PA (PPA) gel and patterned using UV light, which passes through a chrome on quartz photomask. Excess ssDNA is washed from the gel, and cells with membranes displaying lipid-conjugated, complementary ssDNA pattern on PA substrate-bound ssDNA. (**B**) Plot of measured SYBR Gold-labeled 'F' ssDNA spot diameters patterned on PPA substrates vs. nominal diameter of corresponding photomask features (circular spots).

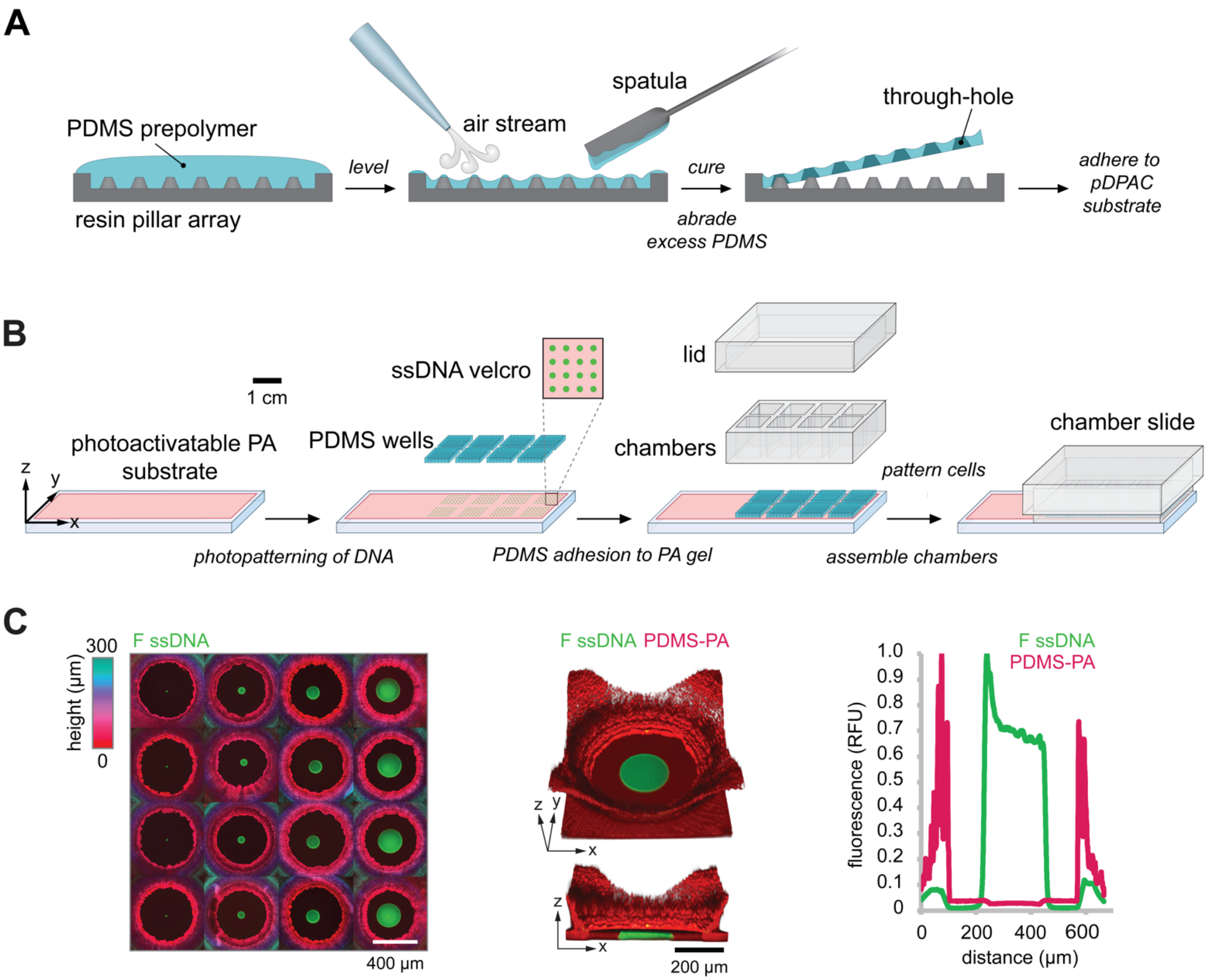


**Fig. S3: Addition of microwell walls and culture chambers for long-term organoid culture.** (**A**) Fabrication of PDMS through-hole sheets by replica micromolding. Resin pillar arrays were 3D-printed using low force stereolithography and then used as a replica mold for PDMS silicone casting. PDMS pre-polymer is poured on a pillar array and leveled with a flat-edged spatula. A stream of compressed air breaks the connection between the pre-polymer residing on top of the pillars and that which is drawn up the sides of the pillars by capillary action. The PDMS is then cured, discs of silicone are abraded off the tops of pillars, and the molded through-hole sheet is removed from the resin substrate. (**B**) Schematic of PA gel cell patterning substrate photopatterned with adhesive ssDNA, non-adhesive PDMS microwell overlays, and their assembly in standard chamber slide format for 8-plex microwell cultures. (**C**) *Left*, confocal immunofluorescence micrographs of ssDNA patterns labeled with SYBR Gold. Height of the PDMS microwell array overlay is encoded by color based on confocal imaging of a rhodamine-methacrylamide co-monomer incorporated into the non-adhesive PA coating. *Middle*, 3D rendering of example ssDNA feature and associated microwell. *Right*, fluorescence profiles of ssDNA and PDMS microwell array coating.

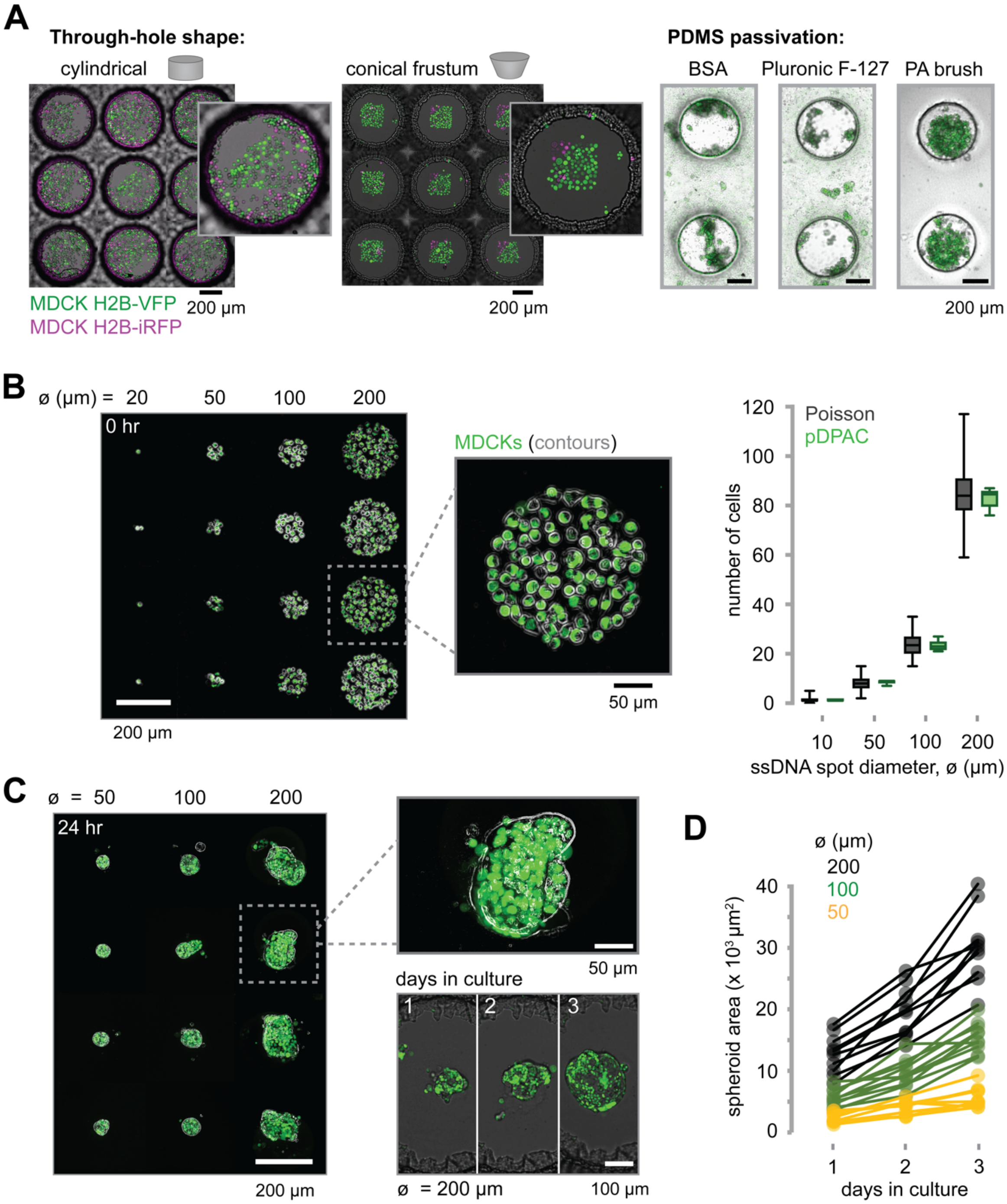


**Fig. S4: Microwell engineering enables precise cell patterning with low non-specific background and adhesion properties.** (**A**) *Left*, conical microwell design reduces non-specific cell patterning. Confocal fluorescence micrographs of typical patterning substrate appearance after performing pDPAC in microwells with PDMS walls having cylindrical or conical profiles. Incomplete and variable washout of non-specifically adhered cells is typically observed for cylindrical wells. *Right*, polyacrylamide brush derivatization of PDMS microwell walls enables robust and long-term blocking of cell adhesion. Confocal fluorescence micrographs of microwells blocked with bovine serum albumin (BSA), Pluronic F-127, or linear polyacrylamide (PA brush), passively seeded with MDCK H2B-VFP cells, and cultured for 3 days. (**B**) *Left*, fluorescence micrographs of example MDCK cell patterns over a range in ssDNA spot sizes. *Right*, plot of patterned cell numbers by spot diameter (mean ± S.D., $n \geq 9$ spots per condition), along with Poisson distribution expected for passive microwell seeding with $\lambda$ = mean of experiment distribution, and Poisson SD = $\sqrt{\lambda}$. (**C**) *Left*, corresponding MDCK spheroids at 72 hrs after patterning in 2D. *Right*, micrographs of an example organoid from a 200-µm ssDNA pattern over the course of three days in culture. (**D**) Growth curves for representative spheroids in each ø group ($n$ = 10 organoids per ø).

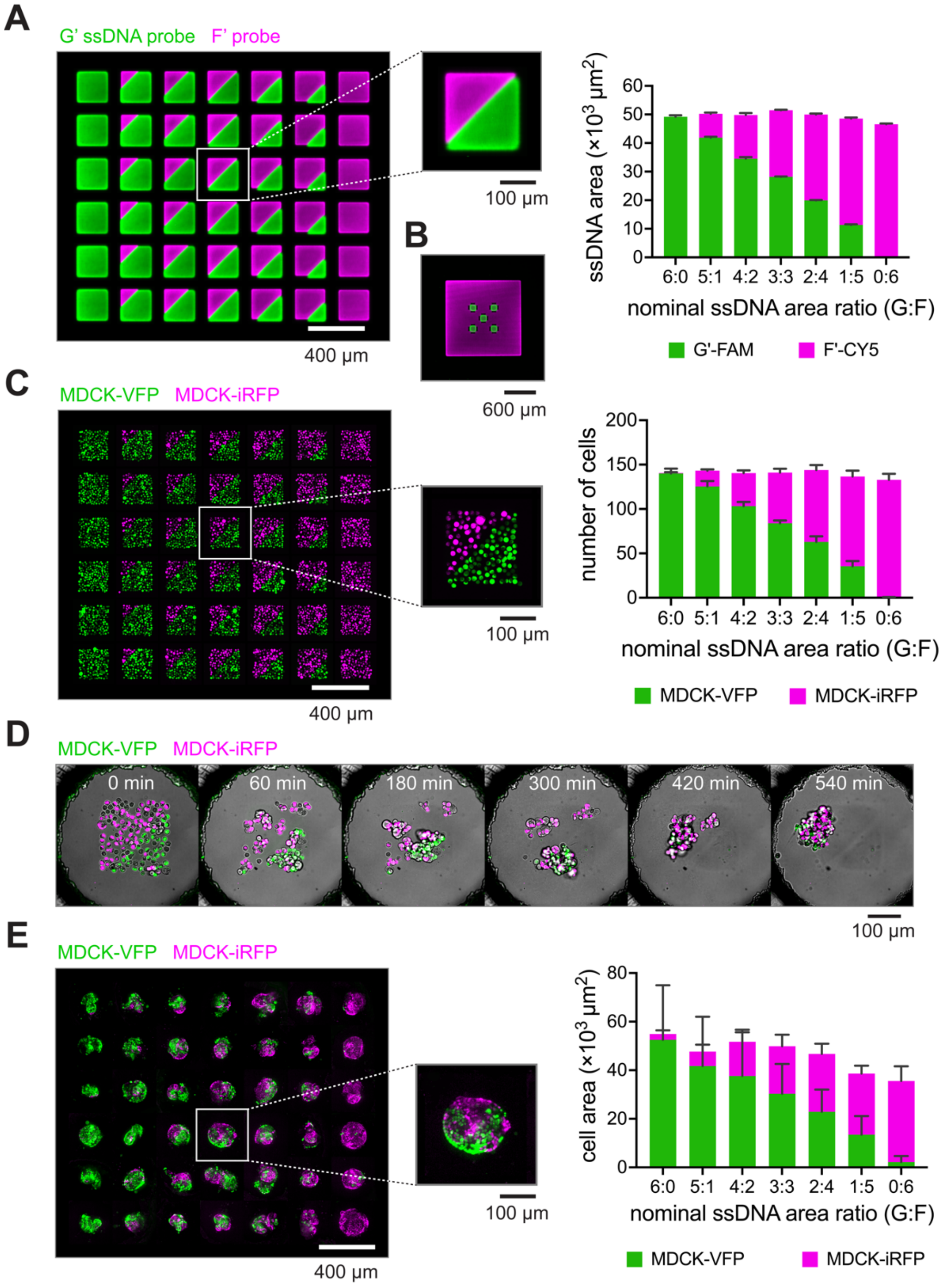


**Fig. S5: Precise cell number and ratio control is maintained after inducing a 2D-to-3D transition in culture context.** (**A**) *Left*, confocal fluorescence micrograph montage of ssDNA patterns over a range of nominal G and F ssDNA surface area ratios concatenated from 42 example microwells. *Right*, histogram of measured pattern areas (mean ± S.D., $n$ = 6 patterns per area ratio) against nominal area ratios (measured from the photomask). (**B**) Confocal immunofluorescence micrograph of successive ssDNA fiducial patterns on a pDPAC substrate. G ssDNA is patterned first and stained with a G'-FITC probe, creating a cross-hair that is positionally aligned with the next photomask for F ssDNA patterning. Simultaneous alignment of similar marks positioned elsewhere on the mask enables rotational alignment. Here, F ssDNA is stained with a F'-CY5 probe. (**C**) Corresponding micrographs and cell number histogram after MDCK cell patterning (mean ± S.D., $n$ = 10 patterns per area ratio). (**D**) Confocal fluorescence micrographs taken from a time-lapse of mosaic spheroid condensation. (**E**) *Left*, sum slices projection micrograph montage of condensed 3D mosaic spheroids created from the 2D MDCK cell patterns in (**C**) at 72 hours in culture. *Right*, histogram of sum of H2B-FP marker areas in 10-μm step confocal planes over the range of ssDNA patterning strand area ratios (mean ± S.D., $n$ ≥ 6 patterns per area ratio).

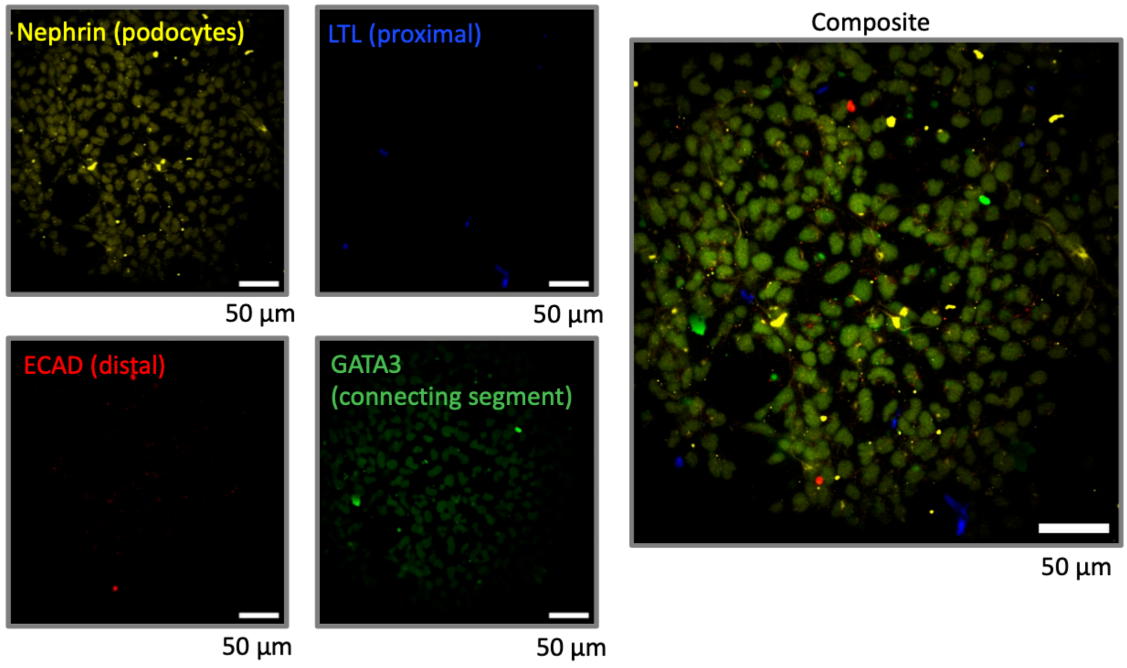


**Fig. S6: Patterned nephron progenitors do not yet express markers of early nephron cell lineages.** Immunofluorescence micrographs of day 10 hiPSC-derived nephron progenitors assayed for expression of the indicated early nephron markers for podocyte, proximal, distal, and connecting segment lineages. All fluorescence profiles are consistent with negative staining.

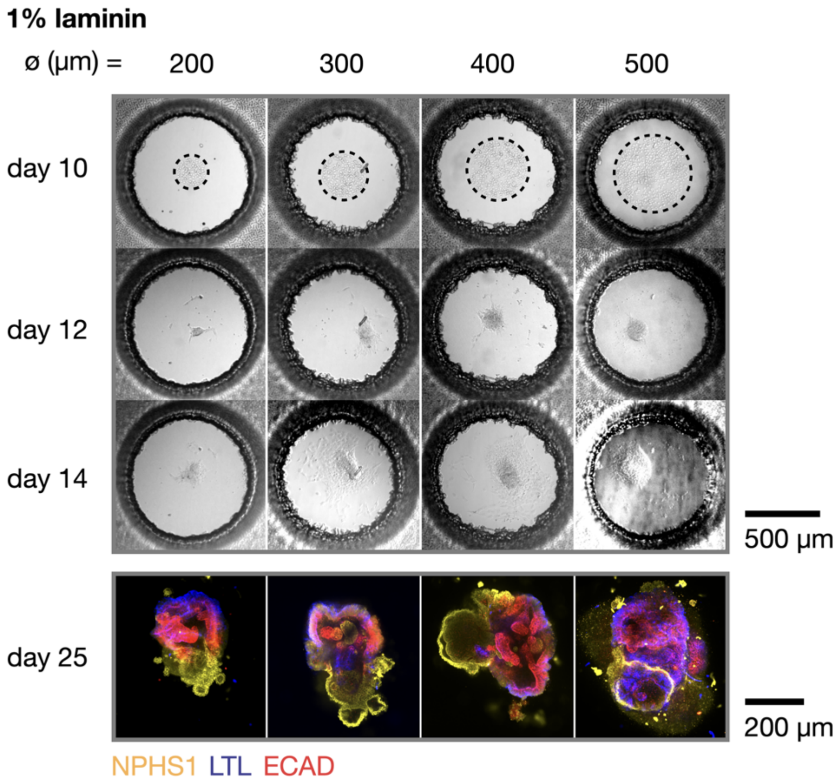


**Fig. S7: Similar aggregation and size-controlled organoid formation for human recombinant laminin culture conditions.** Brightfield and immunofluorescence confocal micrographs during differentiation of nephron progenitor organoids in 1% laminin media after patterning on circular ssDNA patterns with the indicated diameter ø. Dotted lines emphasize the extent of 2D cell patterns prior to transition to 3D culture.

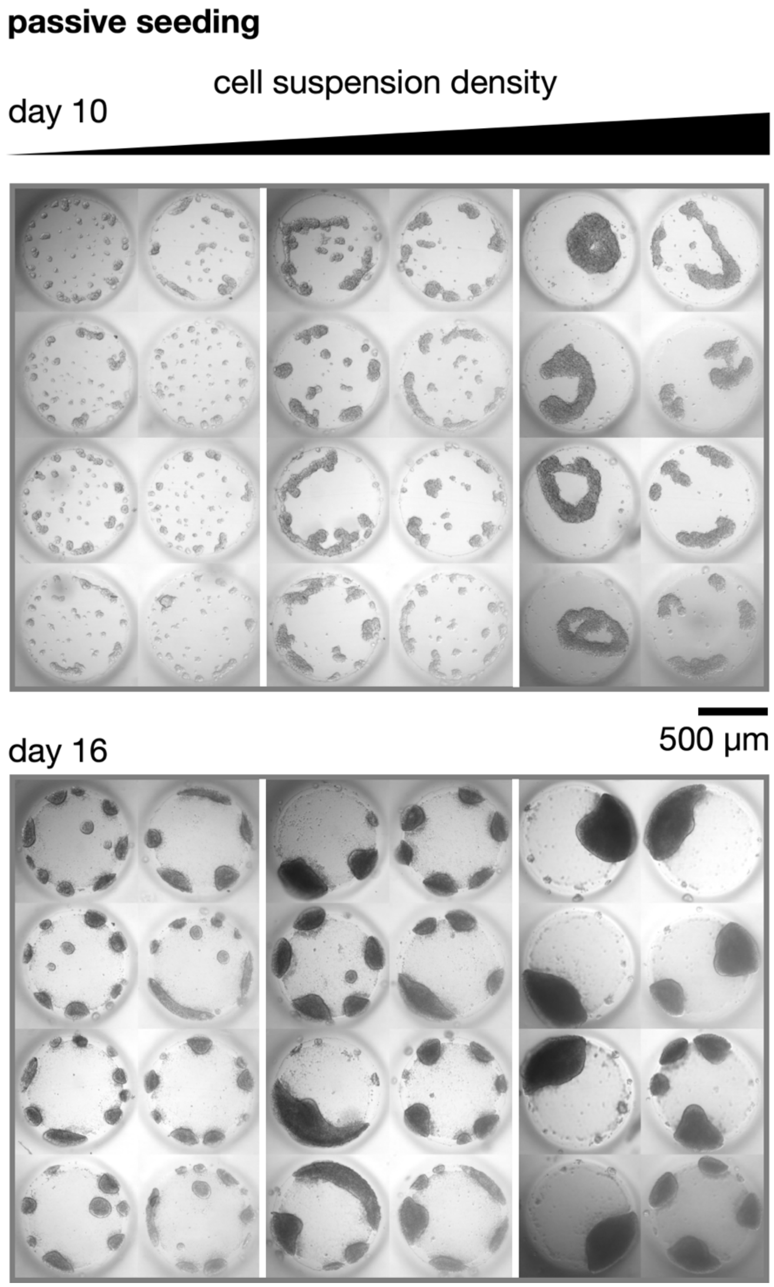


**Fig. S8: Passive nephron progenitor seeding rather than cell patterning leads to unpredictable organoid number and size.** Brightfield confocal micrographs of microwells lacking ssDNA patterns seeded with nephron progenitors over a range of cell densities (0.5, 1, 2x$10^5$ cells per chamber of an 8-chambered slide) and imaged 2 hours after seeding and 6 days later.

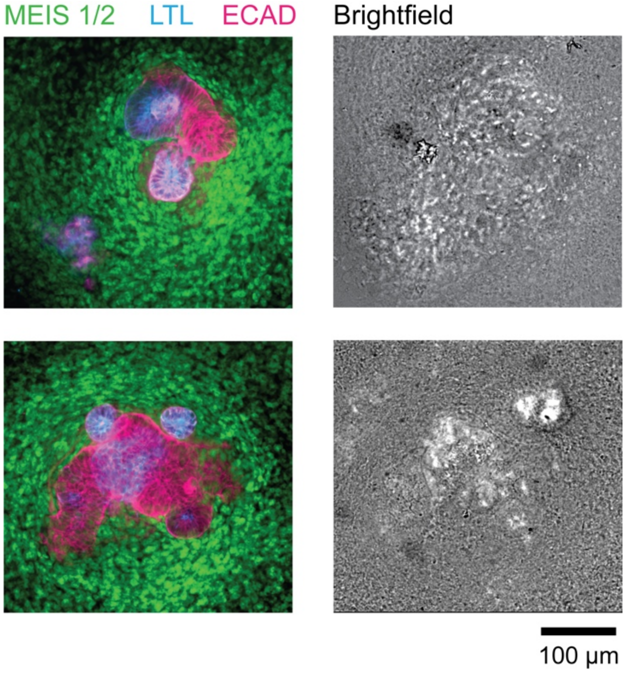


**Fig. S9: Stromal-like cells surround epithelial structures in organoids.** *Left*, confocal micrographs of nephron organoids, stained for markers of proximal tubule (LTL), distal tubule (ECAD+ LTL-), and stroma (MEIS 1/2). *Right*, corresponding brightfield confocal micrographs.

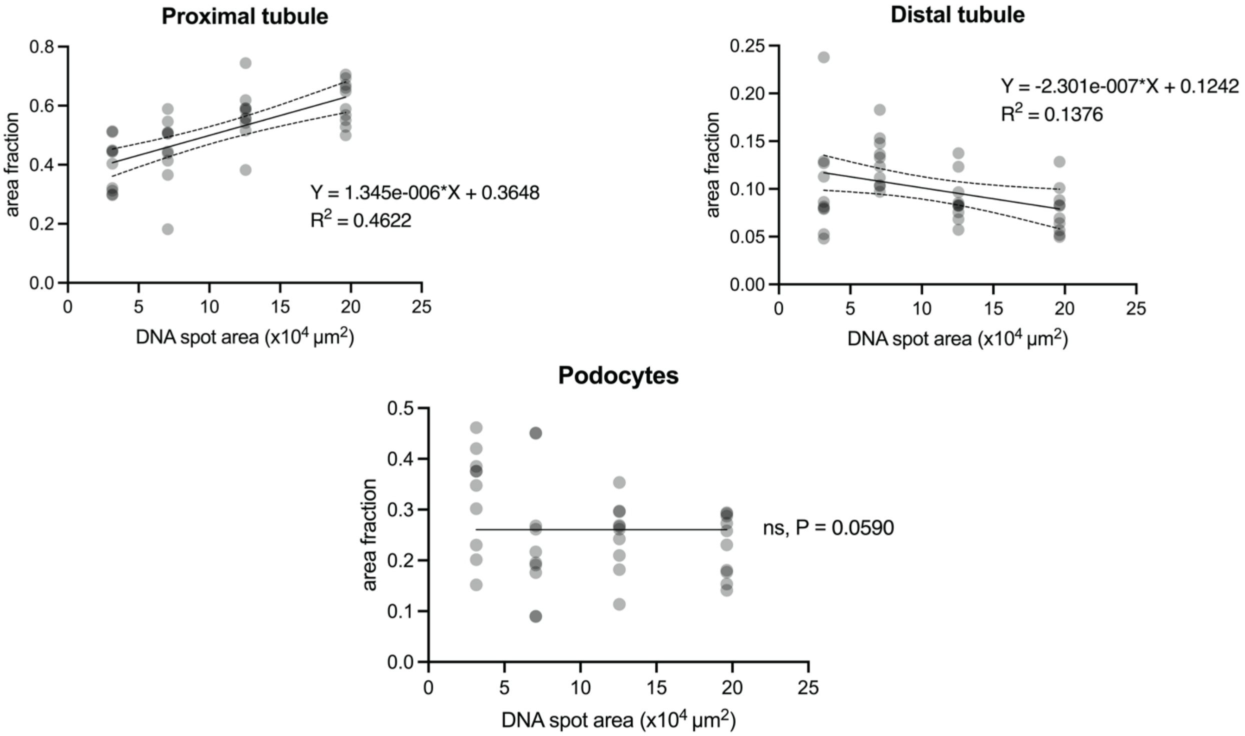


**Fig. S10: Trends in the emergence of different cell types along the early nephron proximal-distal axis, dependent upon the initial numbers of nephron progenitors.** Results of curve fitting using least squares linear and nonlinear regression methods of proximal tubule, distal tubule, and podocyte tissue proportions in organoids at 15 days post cell patterning. Both proximal and distal tubule tissue segmentation data have sloped lines of best fit with statistical significance from zero (proximal $p < 0.0001$, distal $p = 0.0185$), whereas podocyte data fits better to a horizontal line ($n = 10$ organoids per DNA spot size).

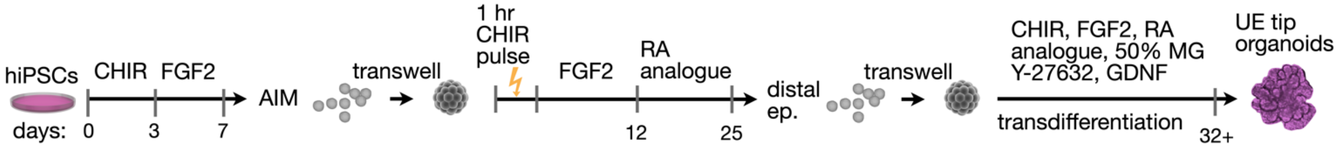


**Fig. S11: Production of hiPSC-derived ureteric epithelium.** Schematic of hiPSC differentiation to anterior intermediate mesoderm, followed by differentiation to distal nephron epithelium and transdifferentiation to ureteric epithelial tip-like cells[1].

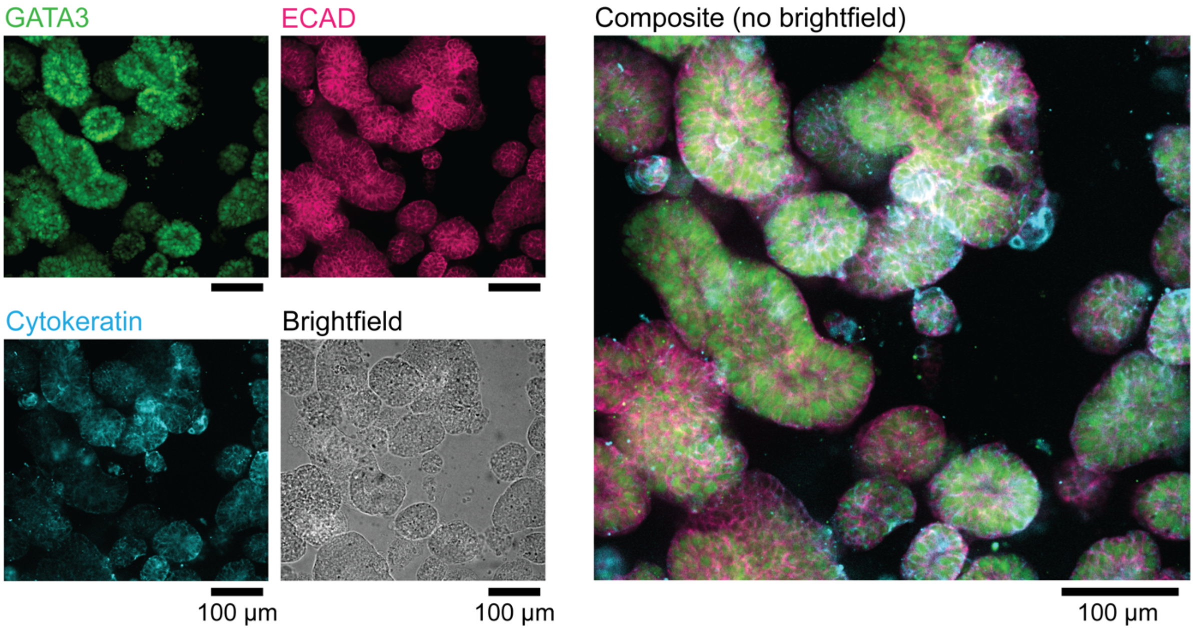


**Fig. S12: UB tip organoids transdifferentiated from hiPSC-derived distal nephron epithelium display markers consistent with the UB tip cell identity.** Brightfield and immunofluorescence confocal micrograph of UB tip organoids, stained for GATA3 (green), E-cadherin (bright pink), and cytokeratin (cyan).

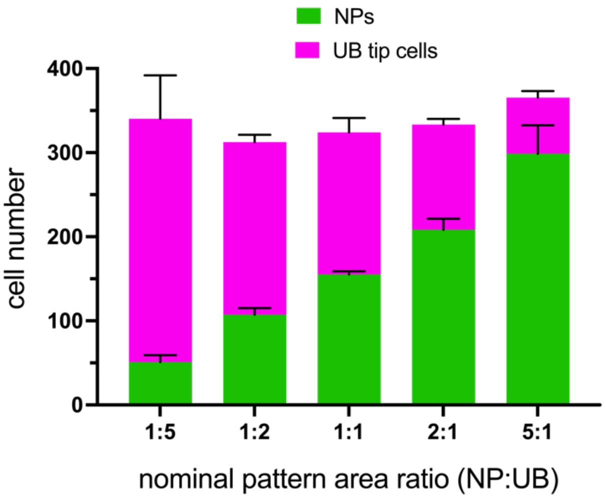


**Fig. S13: Dual NP and UB tip cell patterning closely matches nominal ssDNA area ratios.** Histogram of NPs and UB tip cells patterned on square, 300-µm ssDNA patterns at 5 different nominal area ratios. NPs were patterned on F ssDNA and UB tip cells were patterned on G ssDNA (mean ± S.D., $n \geq 4$ patterns per area ratio).

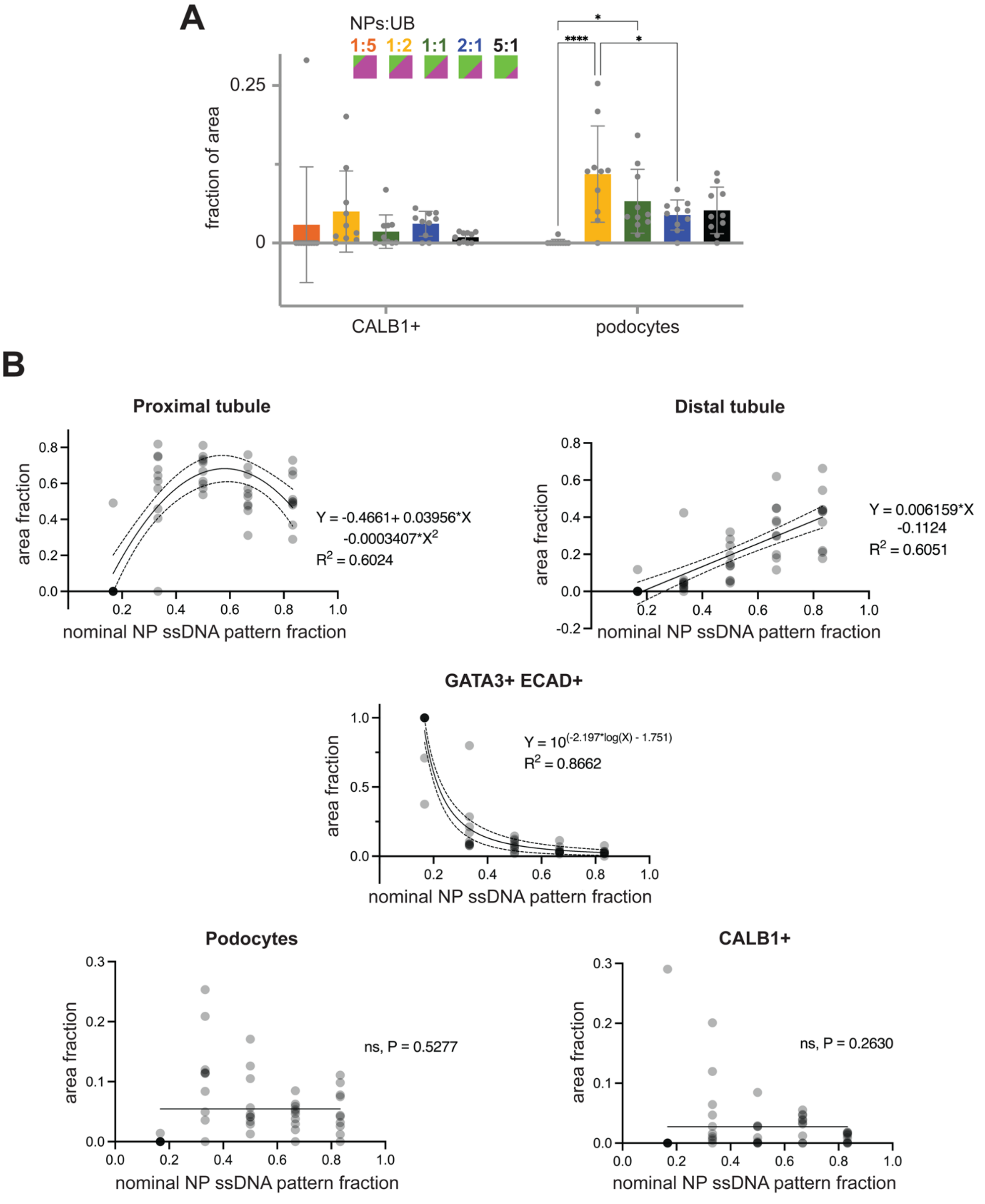


**Fig. S14: Trends in the emergence of different tissue types in mosaic NP/UB tip cell organoids, dependent upon the initial ratios of progenitors.** (**A**) Plot of mosaic organoid CALB1+ and NPHS1+ podocyte tissues (ratio of cell type area to total area of all cell types measured) for organoids formed from 2D co-patterns of different NP:UB tip cell ratios (± S.D., 1-8 slices per *n* = 10 organoids per ratio, Tukey's multiple comparisons test, *p < 0.0332, **p < 0.0021, ***p < 0.0002, ****p < 0.0001). (**B**) Results of curve fitting using least squares linear and nonlinear regression methods for proximal tubule, distal tubule, GATA3+ ECAD+ tissue, podocyte tissue, and CALB1+ tissue proportions in mosaic organoids at 16 days post cell patterning. Proximal tubule, distal tubule, and GATA3+ ECAD+ tissue segmentation data have lines of best fit to the provided models with statistical significance from zero (proximal p = 0.0003, distal p < 0.0001, GATA3+ ECAD+ p < 0.0001), whereas podocyte and CALB1+ data fit better to a horizontal line (*n* = 10 organoids per ratio pattern).

**Supplemental notes**

*Supplemental Note 1: Adhesion and alignment of PDMS through-hole overlays on PA base gels patterned with ssDNA.* When binding PDMS to a substrate, both the PDMS and substrate are typically oxygen plasma-treated, creating reactive groups on both surfaces that create covalent bonds[2], which may disrupt ssDNA integrity, necessitating an alternate method. Instead, we plasma treated only the polyacrylamide brush layer grafted on the PDMS surface before adhering it to the ssDNA-patterned PA substrate. Plasma treatment creates reactive amide groups on the brush layer that hydrogen bond with the PA substrate[3]. For alignment, we used DI water for two reasons: 1) it allowed us to float through-hole sheets above the ssDNA patterns (visualized using SYBR™ Gold staining) and manually align their positions before pressing them into tight contact with the substrate, and 2) we hypothesize that the swelling of the polyacrylamide substrate and brush layer increased interfacial entanglement of the polymer chains, improving adhesion[4]. Once the water had evaporated, the through-hole sheets were immobilized on the polyacrylamide substrate. We then used a thermal bonding process at 80°C to further improve adhesion of the PDMS overlay to the ssDNA-patterned substrate[2]. Thermal annealing did not damage ssDNA patterns, as dry DNA remains stable at temperatures below 130°C[5].

*Supplemental Note 2: Reducing nonspecific cell adhesion in microwells.* One major design challenge was to ensure efficient cell capture on ssDNA within microwells, while discouraging non-specific capture of cells. We found significant advantages here by using a conical rather than a cylindrical well profile, aiding in wash-out of unpatterned cells. Secondly, we required an extremely low cell attachment profile for the microwell walls in order to prevent initial nonspecific cell adhesion as well as organoid spreading and migration out of microwells over relatively long differentiation times. Despite its biofouling property, PDMS was an attractive material for microwell wall fabrication because it is chemically inert, mechanically stable, biocompatible, inexpensive, and easily moldable[6]. We found that among other passive blocking schemes, grafting a non-adhesive linear polyacrylamide brush layer onto the PDMS surface gave the most favorable nonadhesive properties[2,7–9] (**Fig. S4**).

*Supplemental Note 3: Exogenous canonical Wnt activation in mosaic NP/UB tip cell organoids*. We found that applying a CHIR pulse to NPs prior to lifting cells had detrimental effects on NP patterning efficiency within microwells, as it promoted clumping of cells in suspension and nonspecific cell adhesion to microwell walls. We therefore patterned both NPs and UB tip cells and supplied the 1-hr CHIR pulse 24 hrs later on a background of FGF9 treatment, during which time the mosaic organoids were condensing. We found that this CHIR pulse was necessary for NP-derived nephron structures to form by day 26. This contrasts with recent results by Howden et al.[1], where the presence of UB tip cells alone was sufficient to induce nephrogenesis from NPs in a bulk co-culture setting without the addition of small molecules or growth factors. This may have resulted from a smaller ‘community effect’ due to the significantly lower cell mass in our mosaic micro-organoids compared to bulk co-cultures.

**Supplementary Movies**

**Movie S1. Conical microwell integrated with patterned ssDNA.** 3D rendering of confocal z-stack, showing rhodamine-methacrylamide co-monomer (red) incorporated into basal PPA gel and non-adhesive PA coating on conical PDMS walls (not visible). The patterned F ssDNA spot on the PPA substrate is labeled with SYBR Gold (green).

**Movie S2. Condensation of mosaic MDCK spheroids.** Confocal time-lapse of CellTracker-labeled MDCK cell populations condensing into a 3D spheroid after patterning in 2D by pDPAC.

**Movie S3. Condensation of hiPSC-derived nephron progenitor cells.** Brightfield confocal time-lapse showing NPs patterned in 2D by pDPAC rapidly condensing into a 3D spheroid after cleaving dsDNA adhesions to the substrate using DNase (right) vs control without addition of DNase (left).
